# Polyacrylamide-based Antimicrobial Copolymers to Replace or Rescue Antibiotics

**DOI:** 10.1101/2024.09.13.612993

**Authors:** Shoshana C. Williams, Madeline B. Chosy, Carolyn K. Jons, Changxin Dong, Alexander N. Prossnitz, Xinyu Liu, Hector Lopez Herandez, Lynette Cegelski, Eric A. Appel

**Author notes:** **Corresponding author** Eric Appel.

## Abstract

Antibiotics save countless lives each year and have dramatically improved human health outcomes since their introduction in the 20^th^ century. Unfortunately, bacteria are now developing resistance to antibiotics at an alarming rate, with many new strains of “superbugs” showing simultaneous resistance to multiple classes of antibiotics. To mitigate the global burden of antimicrobial resistance, we must develop new antibiotics that are broadly effective, safe, and highly stable for the purpose of global access. In this manuscript, we report the development of polyacrylamide-based copolymers as a novel class of broad-spectrum antibiotics with efficacy against several critical pathogens. We demonstrate that these copolymer drugs are selective for bacteria over mammalian cells, indicating a favorable safety profile. We show that they kill bacteria through a membrane disruption mechanism, which allows them to overcome traditional mechanisms of antimicrobial resistance. Finally, we demonstrate their ability to rehabilitate an existing small-molecule antibiotic that is highly subject to resistance development by improving its potency and eliminating the development of resistance in a combination treatment. This work represents a significant step towards combatting antimicrobial resistance.

## INTRODUCTION

Antimicrobial resistance (AMR) is a growing global crisis. It is estimated that by 2050, AMR will cause over 10 million deaths annually.^1^ In the US alone, AMR already causes over 35,000 deaths annually and costs over $4.6 billion in direct healthcare spending.^2,3^ Unfortunately, progress in the development of new antibiotics has waned, while AMR bacteria arise at an alarming rate.^4,5^ Even as the identification of multidrug resistant (MDR) isolates increases,^6^ publications on antibiotics are decreasing (Figure 1A).^7^

**Figure 1.**
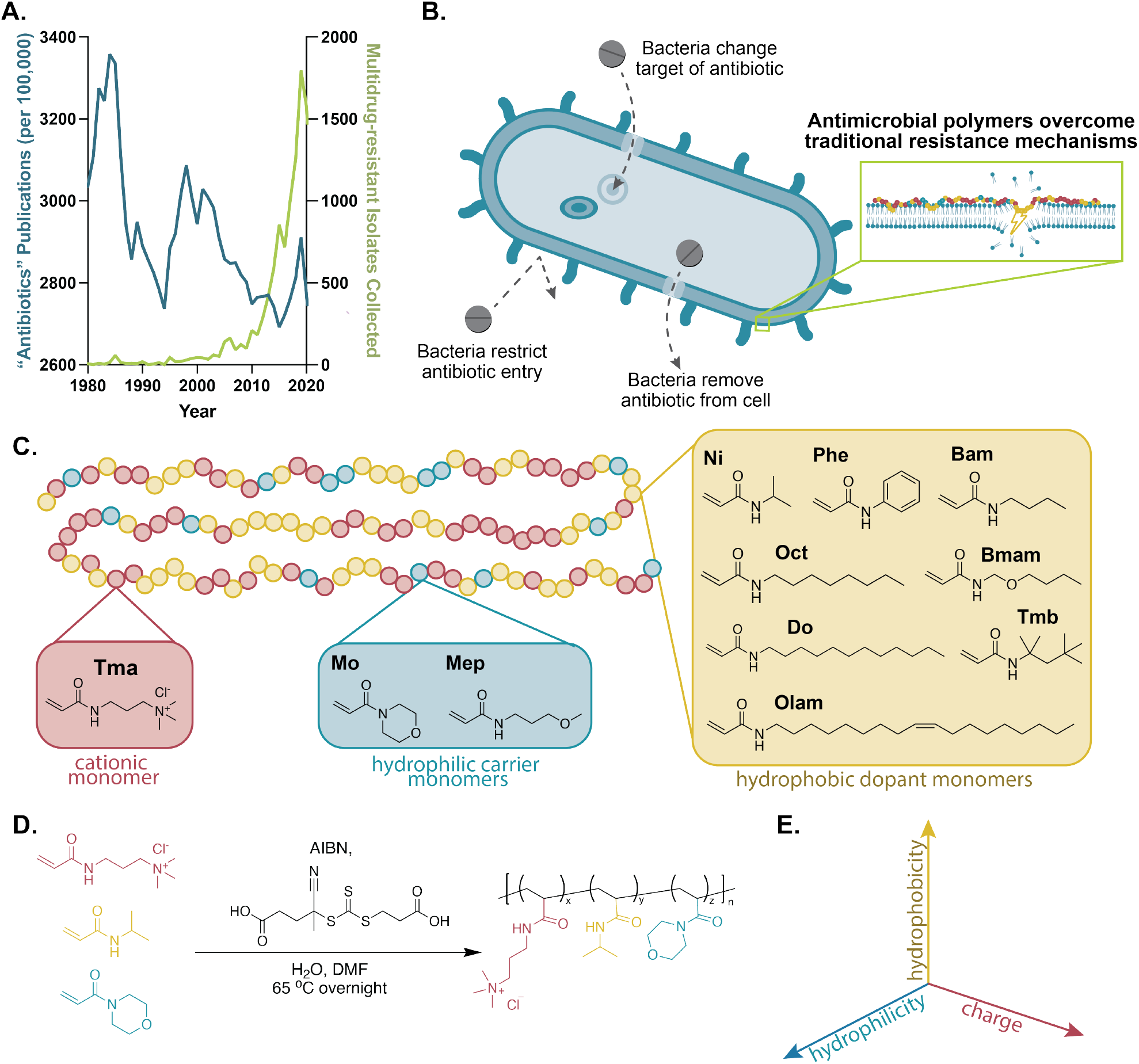
Novel polymers to combat antimicrobial resistance. **A**. Graph comparing the decreasing research into antibiotics with the increasing incidence of MDR bacteria. **B**. Schematic showing common mechanisms of AMR and the hypothesized ability of copolymers to overcome these mechanisms. **C**. Schematic of statistical copolymer library showing monomers used and their classes. **D**. RAFT polymerization conditions. **E**. The use of a ternary system allows for independent tuning of charge and hydrophobicity.

The mechanisms of AMR are molecularly defined and include inhibition of drug uptake, modification of the drug target, inactivation of the drug, and enhanced drug efflux (Figure 1B).^8,9^ Novel antibiotics that could evade these mechanisms are of immense interest to combat the growing threat of AMR.

Previous work has explored the development of antimicrobial polymers and oligomers, both as coatings for devices or surfaces^10–18^ and as treatments.^19–48^ These studies have identified positive charge and hydrophobicity as key parameters to enable antibacterial activity, and several of these polymers were shown to work through a membrane disruption mechanism.^49–62^ This mode of action overcomes traditional resistance mechanisms, since the polymers don’t need access to the intracellular space, and they aren’t specific to a singular molecular target where mutations confer resistance.^63^ Despite these exciting developments, the further translation of these polymers has been hampered by challenging synthesis and toxicity by hemolytic activity. Several approaches have relied on multi-step syntheses to prepare monomers and perform post-polymerization modifications, which can be costly and time-intensive.^29,30,35–38,45,46,62^ Other investigations have utilized sequence-controlled polymers, which similarly require greater synthetic control.^28,31–34,64^ Even with these approaches, several candidates have shown unfavorable hemolytic activity and toxicity profiles.^29,33–36^ To address these challenges, we generated a library of polyacrylamide-derived copolymers for use as antimicrobial agents. Polyacrylamides were selected for their impressive chemical stability, commercial availability, and ease of synthesis. We investigated their activity, safety, mechanism of action, and ability to rescue the effectiveness of existing small-molecule antibiotics.

## RESULTS AND DISCUSSION

### Development of a library of polyacrylamide derivative copolymers

We developed a library of unique polyacrylamide derivatives designed to arrest bacterial growth through a membrane disruption mechanism, which doesn’t rely on specific protein transporters or pathways subject to resistance mechanisms. Each copolymer comprises various ratios of three monomers, including: (i) a cationic monomer to drive polymer adsorption to the negatively charged surface of bacteria, (ii) a hydrophilic “carrier” monomer to tune water solubility, and (iii) a hydrophobic “dopant” monomer to disrupt the bacterial membrane (Figure 1C). We designed this library to vary monomer identity, charge density, hydrophobicity, and molecular weight to allow us to explore how these parameters affect activity and safety. To this end, the introduction of the hydrophilic carrier monomer was significant, not only for its ability to impart water solubility, but also for its utility in allowing for the independent variation of charge and hydrophobicity (Figure 1E). Previous work in the field has explored these parameters through the development of polymers where the hydrophobicity and charge were linked, either in the same monomers or using two monomers.^10,28,35,36^ Inclusion of a third monomer allowed for greater control in exploring the independent effects of charge and hydrophobicity. It also allowed for control of hydrophilicity, which is known to impact antimicrobial activity.^65^

Copolymers were synthesized through statistical copolymerization using a variety of commercially available or easily synthesized monomers by reversible addition-fragmentation chain transfer (RAFT) polymerization (Figure 1D). The copolymers were analyzed by NMR (Figure S3) and GPC (Figure S4). Molecular weights were determined by comparison to PEG standards, except for L-Tmb_5_Mo_90_. Because of this formulation’s low cationic density and higher hydrophobicity, it was more appropriately evaluated by comparison to PMMA standards. The dispersities are typical for controlled radical polymerization techniques (Table 1).

**Table 1.** Composition of polyacrylamide library.

|  | Target DP | Target M<sub>w</sub> (kDa) | M<sub>w</sub> (kDa) | Đ | % Conversion | % Yield |
| --- | --- | --- | --- | --- | --- | --- |
| L-Ni<sub>31</sub>Mo<sub>10</sub> | 70 | 11.1 | 11.1 | 1.12 | n.d. | 103 |
| L-Ni<sub>31</sub>Mep<sub>10</sub> | 70 | 11.1 | 11.9 | 1.12 | n.d. | 113 |
| L-Phe<sub>31</sub>Mo<sub>10</sub> | 70 | 12.4 | 10.3 | 1.16 | n.d. | 94.5 |
| L-Phe<sub>31</sub>Mep<sub>10</sub> | 70 | 12.4 | 10.5 | 1.14 | n.d. | 99 |
| L-Do<sub>31</sub>Mo<sub>10</sub> | 70 | 14.4 | 10.4 | 1.36 | n.d. | 92.5 |
| L-Do<sub>31</sub>Mep<sub>10</sub> | 70 | 14.4 | 12.1 | 1.15 | n.d. | 58 |
| L-Ni<sub>13</sub>Mo<sub>4</sub> | 70 | 12.8 | 13.6 | 1.15 | n.d. | 94 |
| L-Ni<sub>13</sub>Mep<sub>4</sub> | 70 | 12.8 | 13.6 | 1.18 | n.d. | 65 |
| L-Phe<sub>13</sub>Mo<sub>4</sub> | 70 | 13.5 | 12.6 | 1.16 | n.d. | 92.5 |
| L-Phe<sub>13</sub>Mep<sub>4</sub> | 70 | 13.5 | 11.5 | 1.25 | n.d. | 81 |
| L-Do<sub>13</sub>Mo<sub>4</sub> | 70 | 14.4 | 11.9 | 1.32 | n.d. | 69.5 |
| L-Do<sub>13</sub>Mep<sub>4</sub> | 70 | 14.4 | 12.1 | 1.37 | n.d. | 78.5 |
| H-Ni<sub>31</sub>Mo<sub>10</sub> | 115 | 18.3 | 21.3 | 1.21 | 100 | 91.5 |
| H-Ni<sub>31</sub>Mep<sub>10</sub> | 115 | 18.3 | 22.0 | 1.15 | 93.6 | 85.5 |
| H-Phe<sub>31</sub>Mo<sub>10</sub> | 115 | 20.3 | 22.5 | 1.18 | 99.3 | 84.5 |
| H-Phe<sub>31</sub>Mep<sub>10</sub> | 115 | 20.3 | 19.1 | 1.15 | 100 | 92.5 |
| H-Do<sub>31</sub>Mo<sub>10</sub> | 115 | 23.6 | 21.0 | 1.27 | 100 | 95 |
| H-Do<sub>31</sub>Mep<sub>10</sub> | 115 | 23.7 | 19.3 | 1.24 | 96.7 | 81.5 |
| L-Bam<sub>31</sub>Mep<sub>10</sub> | 70 | 11.7 | 15.1 | 1.25 | 85.6 | 69 |
| L-Bmam<sub>31</sub>Mep<sub>10</sub> | 70 | 12.7 | 15.4 | 1.27 | 88 | 68 |
| L-Tmb<sub>31</sub>Mep<sub>10</sub> | 70 | 13.3 | 11.3 | 1.28 | 99.3 | 63 |
| L-Oct<sub>31</sub>Mep<sub>10</sub> | 70 | 13.3 | 12.6 | 1.22 | 99 | 51 |
| L-Olam<sub>31</sub>Mep<sub>10</sub> | 70 | 15.5 | 10.3 | 1.27 | 96.7 | 71 |
| L-Do<sub>30</sub>Mep<sub>5</sub> | 70 | 14.75 | 14.0 | 1.23 | n.d. | 59 |
| L-Tmb<sub>5</sub>Mo<sub>90</sub> | 70 | 10.16 | 12.8 | 1.21 | n.d. | 100 |
| L-Oct<sub>5</sub>Mep<sub>5</sub> | 70 | 14.07 | 15.0 | 1.20 | n.d. | 100 |
| L-Phe<sub>15</sub>Mo<sub>30</sub> | 70 | 12.06 | 14.2 | 1.18 | n.d. | 89 |

### Antibiotic efficacy and safety of polyacrylamide copolymers

To evaluate the efficacy of each novel copolymer, its minimum inhibitory concentration (MIC) was determined against a standard Gram-negative strain, *E. coli* ATCC 25922, and a standard Gram-positive strain, *S. aureus* ATCC 29213 (Figure 2A, Table 2). Of the 27 copolymers evaluated, 20 showed activity against at least one of these targets, and 11 showed activity against both, indicating broad-spectrum efficacy. We observed no trade-off between efficacy against *S. aureus* compared to *E. coli*, with several copolymers demonstrating robust activity against both (Figure S5). The observed MIC values were comparable to those of previously reported antimicrobial polymers and licensed antibiotic drug products (Table 2).^28–30,32,35,36,46,51,66–68^

**Table 2.**
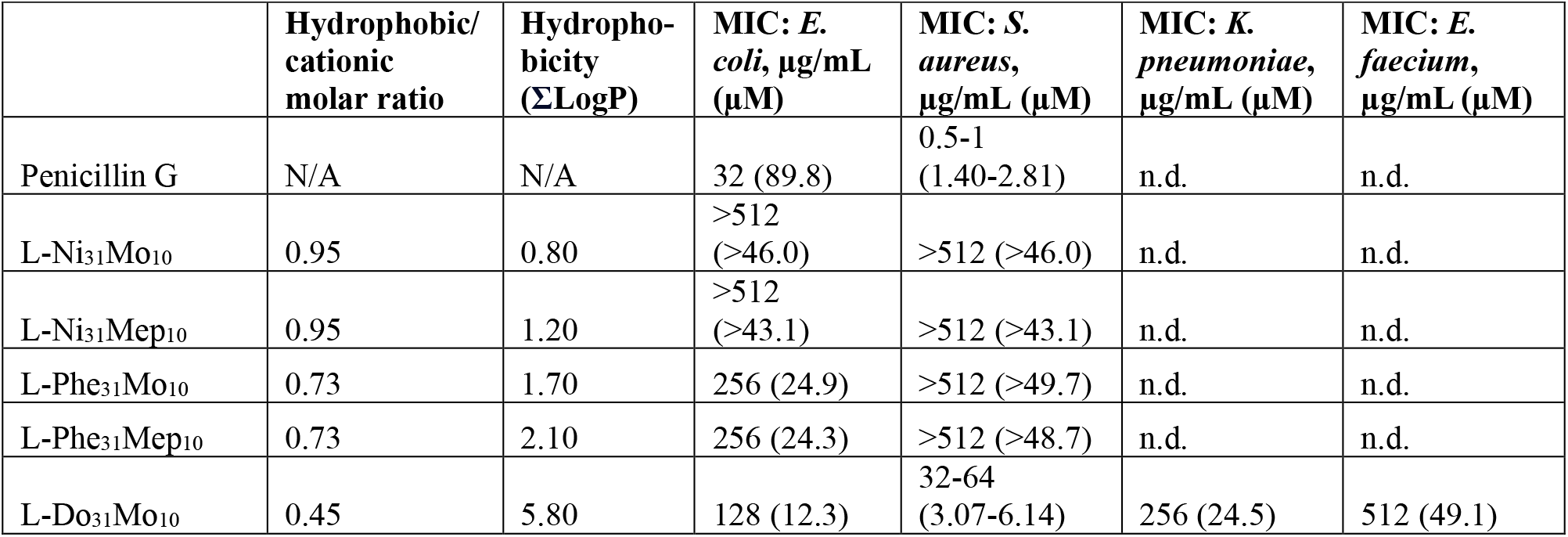
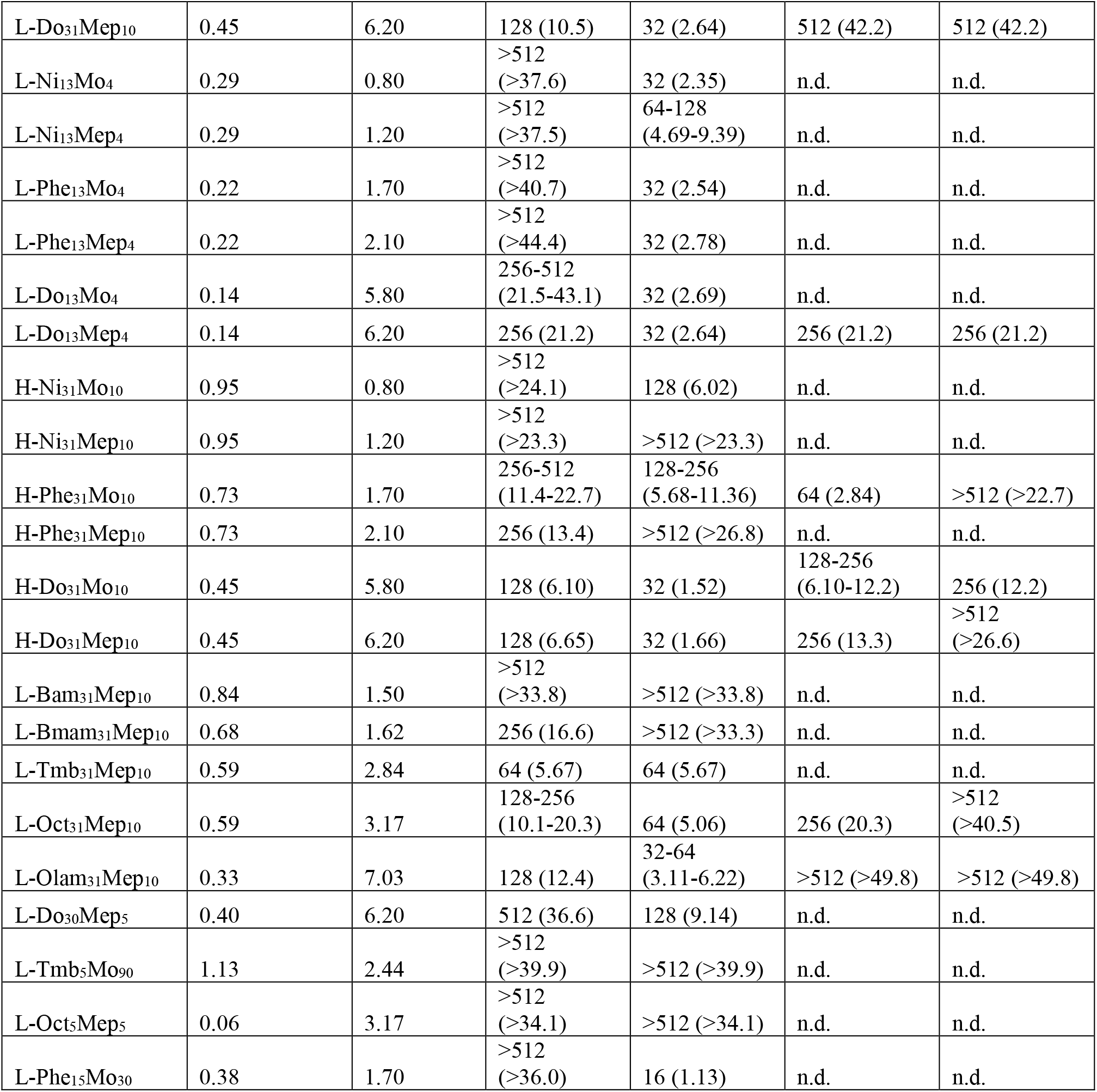
Antibacterial efficacy of novel polyacrylamides and penicillin G.

**Figure 2.**
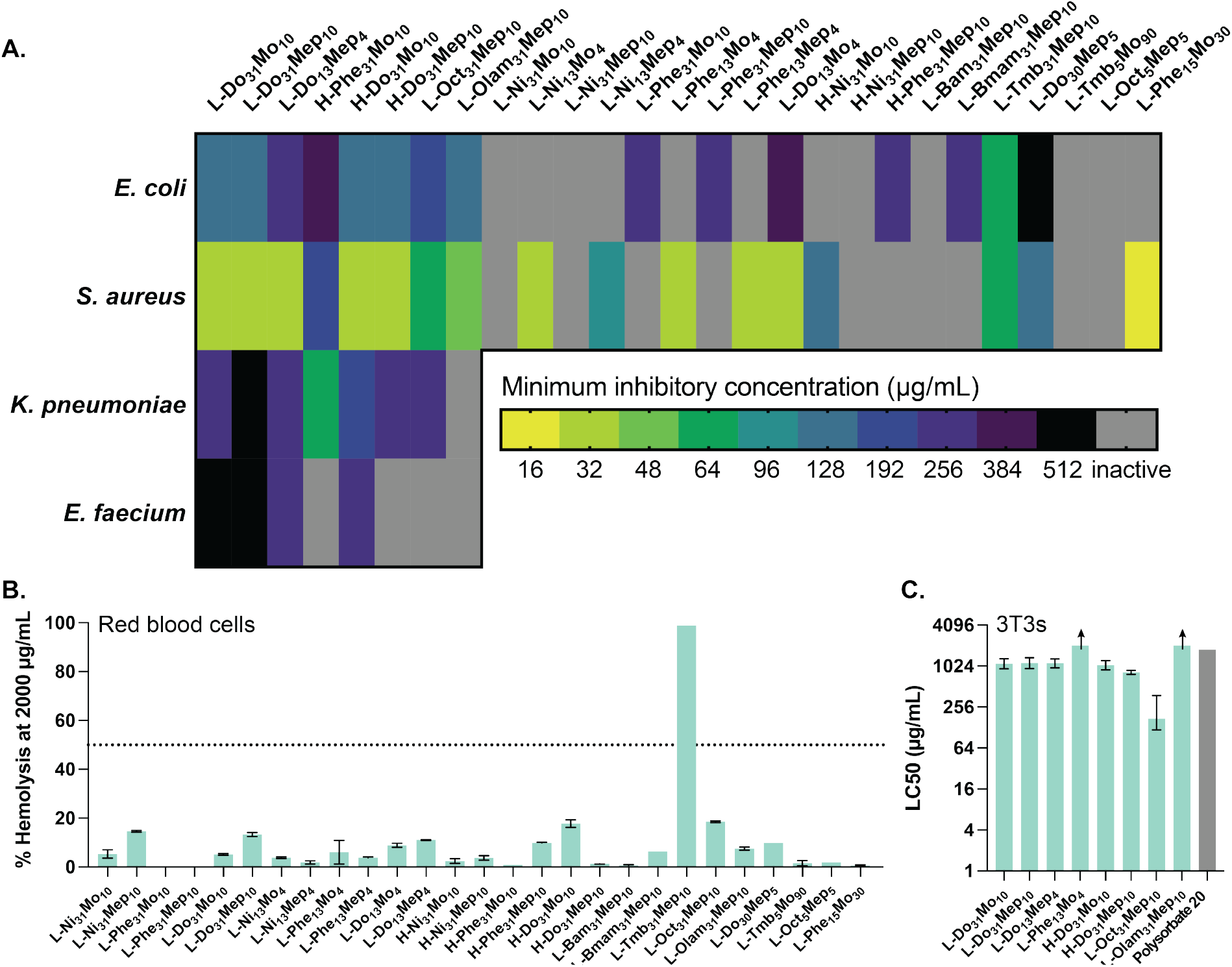
Efficacy and safety of novel polyacrylamides. **A**. Heat map showing the antibacterial efficacy of each polymer against several bacteria. **B**. Hemolytic activity of each polymer at 2000 μg/mL. **C**. LC50 values of eight copolymers against 3T3 cells.

The hemolytic activity of each copolymer was measured as a metric of mammalian non-toxicity. Most copolymers showed remarkably low hemolytic activity, even up to concentrations exceeding 2000 μg/mL, indicating favorable safety profiles (Figure 2B, Table 3). Indeed, for most polymers, HC_50_ (the concentration at which 50% of red blood cells are lysed) values could not be determined because it fell beyond the concentration range tested. Our polymers demonstrated clear selectivity for permeabilizing bacteria, leaving red blood cells intact. Interestingly, the antibacterial efficacy of the copolymer did not directly correlate with the hemolytic activity. The promising safety profile of our copolymers demonstrates the importance of our ternary copolymer design, where the introduction of a hydrophilic monomer enabled access to copolymers with potent antibacterial activity and low toxicity.

**Table 3.** *In vitro* safety of polyacrylamide copolymers and polysorbate 20.

|  | HC<sub>50</sub>, µg/mL<br>(90% CI) | LC50, 3T3s, µg/mL<br>(90% CI) | LC50, 3T3s, µg/mL<br>(90% CI) |
| --- | --- | --- | --- |
| Polysorbate 20 | n.d. | 1800 | 1700 (200-2000) |
| L-Ni<sub>31</sub>Mo<sub>10</sub> | >2000 | n.d. | n.d. |
| L-Ni<sub>31</sub>Mep<sub>10</sub> | >2000 | n.d. | n.d. |
| L-Phe<sub>31</sub>Mo<sub>10</sub> | >2000 | n.d. | n.d. |
| L-Phe<sub>31</sub>Mep<sub>10</sub> | >2000 | n.d. | n.d. |
| L-Do<sub>31</sub>Mo<sub>10</sub> | >2000 | 1100 (900-1300) | 200 (100-300) |
| L-Do<sub>31</sub>Mep<sub>10</sub> | >2000 | 1100 (900-1400) | 80 (40-100) |
| L-Ni<sub>13</sub>Mo<sub>4</sub> | >2000 | n.d. | n.d. |
| L-Ni<sub>13</sub>Mep<sub>4</sub> | >2000 | n.d. | n.d. |
| L-Phe<sub>13</sub>Mo<sub>4</sub> | >2000 | n.d. | n.d. |
| L-Phe<sub>13</sub>Mep<sub>4</sub> | >2000 | n.d. | n.d. |
| L-Do<sub>13</sub>Mo<sub>4</sub> | <50 | n.d. | n.d. |
| L-Do<sub>13</sub>Mep<sub>4</sub> | >2000 | 1100 (1000-1300) | 60 (30-80) |
| H-Ni<sub>31</sub>Mo<sub>10</sub> | >8000 | n.d. | n.d. |
| H-Ni<sub>31</sub>Mep<sub>10</sub> | >8000 | n.d. | n.d. |
| H-Phe<sub>31</sub>Mo<sub>10</sub> | >8000 | >2048 | 200 (90-300) |
| H-Phe<sub>31</sub>Mep<sub>10</sub> | >8000 | n.d. | n.d. |
| H-Do<sub>31</sub>Mo<sub>10</sub> | >8000 | 1100 (900-1200) | 100 (40-300) |
| H-Do<sub>31</sub>Mep<sub>10</sub> | 6000<br>(5000-8000) | 820 (770-880) | 70 (30-110) |
| L-Bam<sub>31</sub>Mep<sub>10</sub> | >8000 | n.d. | n.d. |
| L-Bmam<sub>31</sub>Mep<sub>10</sub> | 6000<br>(5000-8000) | n.d. | n.d. |
| L-Tmb<sub>31</sub>Mep<sub>10</sub> | <62.5 | n.d. | n.d. |
| L-Oct<sub>31</sub>Mep<sub>10</sub> | 5000<br>(4000-6000) | 200 (100-400) | 40 (30-100) |
| L-Olam<sub>31</sub>Mep<sub>10</sub> | >8000 | >2048 | 90 (70-100) |
| L-Do<sub>30</sub>Mep<sub>5</sub> | 3000<br>(3000-4000) | n.d. | n.d. |
| L-Tmb<sub>5</sub>Mo<sub>90</sub> | >4000 | n.d. | n.d. |
| L-Oct<sub>5</sub>Mep<sub>5</sub> | >4000 | n.d. | n.d. |
| L-Phe<sub>15</sub>Mo<sub>30</sub> | >4000 | n.d. | n.d. |

Based on the results of our screen with *E. coli* and *S. aureus*, as well as the hemolysis profiles, eight copolymers were selected as top-performing candidates for further evaluation. These candidates were then tested against two additional bacteria: *K. pneumoniae* ATCC 13884 (Gram-negative) and *E. faecium* ATCC 35667 (Gram-positive). These bacteria, along with *E. coli* and *S. aureus*, are members of a group of virulent pathogens known for their ability to develop resistance to antibiotics.^69,70^ Each of the tested pathogens is on the WHO priority list on account of its threat to global health.^71^ Among these eight copolymers, seven demonstrated activity against at least one additional strain, while four exhibited activity against both. The efficacy of our leading copolymers against these species represents a clear step towards combatting antimicrobial resistance.

To further evaluate the safety of our leading copolymers, we examined the cytotoxicity of our top-performing candidates. We determined the LC50 following 24 h of exposure with 3T3s (Figure 2C, Table 3), and A549s (Figure S6, Table 3). The LC50 values were compared to those for polysorbate 20, a common excipient in FDA-approved drug products. Notably, polysorbate 20 is approved for intranasal administration at a concentration of 25 mg/mL,^72^ well above its LC50 for either of the cell lines tested. The copolymers exhibited lower LC50 values than polysorbate 20 in A549 cells, but more comparable LC50 values to polysorbate 20 in 3T3 cells. These results further demonstrate the safety of these antibiotic copolymer candidates.

### Polyacrylamide copolymer antibiotics function through membrane disruption

Based on the efficacy and safety data, we selected one top candidate copolymer, L-Do_31_Mep_10_, for further characterization. We sought to verify our hypothesized mechanism of cell-killing behavior through membrane disruption. We incubated *E. coli* with the DNA-intercalating dye propidium iodide in the presence of L-Do_31_Mep_10_. We observed a dramatic increase in fluorescence over an hour of monitoring (Figure 3A), indicating that both the outer and inner membranes of *E. coli* had been compromised, thereby allowing the propidium iodide to access the DNA within the bacteria. Our negative control, penicillin G, kills bacteria through inhibiting cell wall synthesis and did not lead to a similar increase in fluorescence.

**Figure 3.**
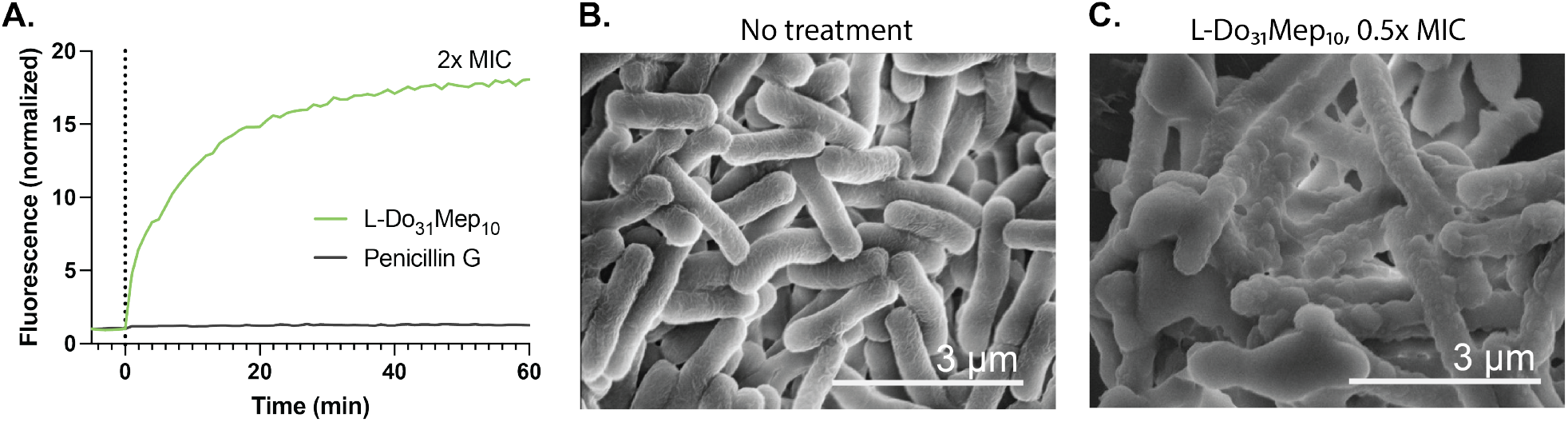
L-Do_31_Mep_10_ disrupts the membrane of *E. coli*. **A**. Membrane permeabilization assay, using the fluorescent probe propidium iodide. **B**. SEM image of *E. coli* (untreated). **C**. SEM image of *E. coli* treated with L-Do_31_Mep_10_.

Further, we visually examined *E. coli* before and after exposure to L-Do_31_Mep_10_ at to half the lethal level (0.5x MIC; 64 μg/mL; 6.27 μM) using scanning electron microscopy (SEM) (Figure 3B-C). Perturbations in the membrane were observed in the presence of L-Do_31_Mep_10_. Notably, blistering was visible in the treated sample, consistent with disruptions to the physical integrity of the membrane.

### Polyacrylamide copolymer antibiotics mitigate onset of resistance

We hypothesized that a membrane disruption mechanism would allow our treatments to overcome traditional mechanisms of resistance. We tested this hypothesis by exposing *E. coli* ATCC 25922 to half the lethal level of L-Do_31_Mep_10_ for 7 passages. In each passage, we calculated the MIC of the bacteria (MIC_n_) and compared it to the MIC of naïve *E. coli* (MIC_0_). We compared the copolymer efficacy to a small molecule control (penicillin G). Penicillin G is not reliably effective against clinical *E. coli* isolates on account of widespread resistance developed over several decades of penicillin G overuse, and newer penicillin derivatives, including ampicillin and amoxicillin, are more commonly used to treat infections in patients.^73^ For these assays we selected a strain of *E. coli* with known susceptibility to penicillin G, in accordance with previous reports.^74,75^ In these studies, no resistance to the copolymer was observed over the course of the study, whereas resistance to penicillin G quickly developed over the same timeframe (Figure 4). These results indicate that the physical membrane disruption mechanism of the copolymer can mitigate the development of resistance.

**Figure 4.**
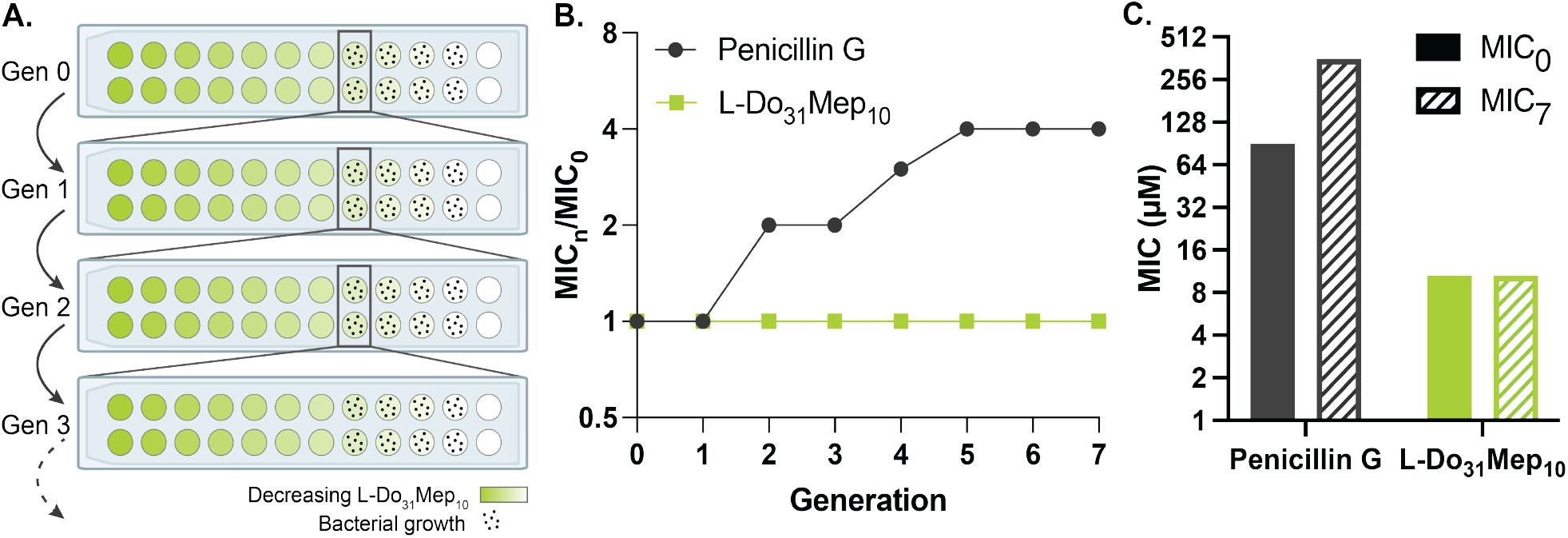
L-Do_31_Mep_10_ evades resistance mechanisms. **A**. Schematic indicating the workflow for the resistance assay. **B**. Results of resistance assay, indicating that L-Do_31_Mep_10_ is immune to resistance over 7 generations. **C**. Comparison of MIC_0_ and MIC_7_ for penicillin G and L-Do_31_Mep_10_.

### Polyacrylamide copolymer antibiotics can rehabilitate traditional antibiotics

We sought to evaluate the potential of our leading polyacrylamide copolymer to improve the efficacy of existing antibiotics. Previous work demonstrated that polymers can improve the delivery and potency of small-molecule antibiotics.^11,13,52^ We hypothesized that, since our novel copolymer disrupts the membrane of bacteria, it could facilitate entry of antibiotics into the cell (Figure 5A). We evaluated a range of antibiotics, including one that is already effective against *E. coli* (rifampicin), one that is ineffective due to an inability to access to the periplasmic space (vancomycin), and one that shows moderate efficacy due to incomplete access to the periplasmic space (penicillin G).^76–78^ We determined the MIC of each antibiotic alone and in the presence of a sublethal amount (0.5x MIC) of L-Do_31_Mep_10_. Cotreatment decreased the MIC by a factor of four for penicillin G and by a factor of two for vancomycin and rifampicin, supporting our hypothesis that the copolymer can improve intracellular access for each of these antibiotics (Figure S7). Thus, clinically relevant antibiotics become more potent in combination treatment with the antibacterial polyacrylamide.

**Figure 5.**
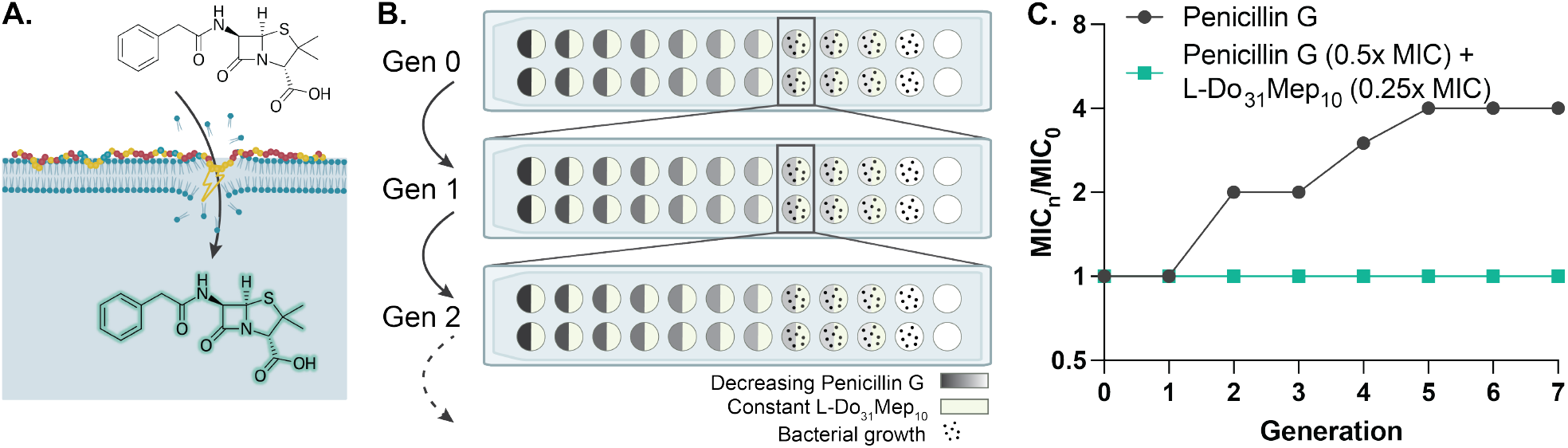
L-Do_31_Mep_10_ improves the efficacy of penicillin G. **A.** Schematic showing the hypothesized mechanism of membrane perturbation that facilitates antibiotic uptake. **B**. Schematic detailing the resistance assay where the copolymer is used as an adjuvant to penicillin G. **C**. Results of resistance assay, indicating that combination treatment prevents resistance against penicillin G over 7 generations.

Additionally, we sought to determine our copolymer’s ability to affect the development of resistance to penicillin G. It has been previously demonstrated that a combination treatment of an antimicrobial polymer and a small-molecule antibiotic can mitigate the development of resistance, although this is likely due to the polymer’s ability to circumvent resistance, as the concentration of the polymer varied alongside the small molecule in this resistance assay.^61^ Another study demonstrated that a polymer could be used to protect a small-molecule antibiotic from the development of resistance; however, the polymer described had minimal antimicrobial activity on its own.^62^ We hypothesized that our antimicrobial polymers, in addition to showing efficacy in isolation, could also act as an adjuvant to protect a small-molecule antibiotic from resistance. To investigate this hypothesis, we repeated the resistance study, but we used L-Do_31_Mep_10_ as an adjuvant at a constant sublethal concentration (0.25x MIC; 32 μg/mL; 2.64 μM). We evaluated the development of resistance to penicillin G over 7 passages in the presence of L-Do_31_Mep_10_, and we observed no resistance (Figure 5B-C). This remarkable finding suggests that our leading copolymer antibiotic may serve as a tool to combat the development of resistance against small-molecule antibiotics that are subject to traditional resistance mechanisms on their own.

### Evaluation of the physicochemical properties driving efficacy of polyacrylamide copolymers

In addition to evaluation of the antibacterial efficacy and safety of our polyacrylamide copolymers, we analyzed each copolymer by some key physicochemical characteristics to investigate the influence of these properties on antibacterial efficacy. We featurized each copolymer using the ChemoPy package to compute hundreds of chemical characteristics.^79^ We condensed these multidimensional data using a principal component analysis (PCA), which preserved 60% of the variation of the dataset (Figure 6A). In this analysis, we noticed little clustering, indicating that our effective copolymers were spread across the chemical space of our library. This observation likely arose because most of our entries were effective against at least one strain of bacteria. An expansion of the library in future studies might yield more robust patterns in the PCA.

**Figure 6.**
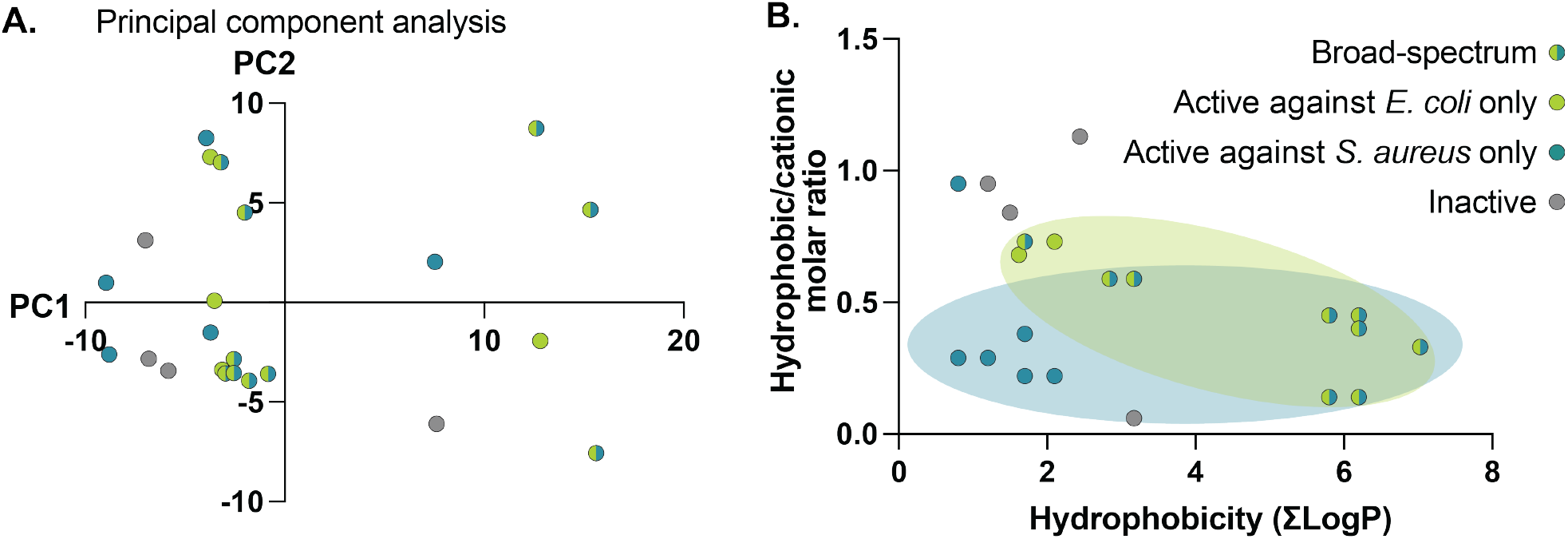
Evaluation of chemical differences in the library of novel polyacrylamides. Each point represents a polymer entry in our library, color-coded by its antimicrobial activity. **A**. Principal component analysis to condense highly dimensional chemical data. **B**. The impact of hydrophobicity and hydrophobic/cationic balance on the efficacy of novel copolymers.

We observed a strong relationship between the hydrophobicity of the monomers used and the antibacterial efficacy of the polymer. To explore this, we calculated the sum of the estimated LogP values (calculated by ChemDraw) for the hydrophobic and hydrophilic monomers used for each polymer (Table 2, Figure 6). We also investigated the influence of the molar ratio of the hydrophobic monomer to the hydrophilic monomer (Table 2, Figure 6). We found that the copolymers with the great breadth of activity were those with the highest LogP values. In contrast, the least effective copolymers had low LogPs and high hydrophobic/cationic molar ratios, suggesting that these copolymers comprised a high concentration of insufficiently hydrophobic monomers. In this analysis, we also noticed that copolymers with a higher charge density showed preferential activity towards *S. aureus*, while those with a slightly higher hydrophobic/cationic ratio were relatively more active towards *E. coli*.

## CONCLUSION

We synthesized a library of 27 polyacrylamide copolymers at gram-scale with high yield using RAFT polymerization. We utilized statistical terpolymers to provide for flexibility in independently varying the charge density and hydrophobicity of our library entries. We evaluated their activity as antibacterial agents against *E. coli, S. aureus, K. pneumoniae*, and *E. faecium* and found several copolymers that showed broad-spectrum activity. Further, we evaluated the safety of these copolymers *in vitro*, and we determined that they were highly selective for bacteria over mammalian cells, leaving the latter intact even at high concentrations. Their safety profiles were comparable to used inactive excipients widely used in FDA-approved drug products.

One candidate copolymer, L-Do_31_Mep_10_, was selected for further characterization on account of its exceptional breadth and safety profile. Imaging studies confirmed that the copolymer functions through a membrane-disruption mechanism of cell-killing, which was found to both enhance the efficacy of traditional small-molecule antibiotics by improving access into the bacteria, as well as to mitigate traditional resistance mechanisms. Furthermore, a combination treatment of this leading copolymer with penicillin G prevented *E. coli* from developing resistance to the penicillin G, despite the copolymer being present well below its own lethal dose. To our knowledge, this is the first report of an antimicrobial copolymer protecting a small molecule from the development of resistance. This remarkable result suggests that these polyacrylamide copolymers may be useful as adjuvants to rehabilitate clinically used antibiotics in the fight against antimicrobial resistance.

## ASSOCIATED CONTENT

The following file is available free of charge:

Supporting Information: experimental methods and supporting data (PDF)

## AUTHOR INFORMATION

## Author contributions

The study was conceptualized by S.C.W. and E.A.A. with guidance from L.C. Experiments were conducted by S.C.W., with contributions from M.B.C, X.L., C.K.J., C.D., and A.N.P. H.L.H. contributed to the principal component analysis. The original draft was written by S.C.W.

## ACKNOWLEDGEMENTS

Part of this work was performed at the Stanford Nano Shared Facilities (SNSF), supported by the National Science Foundation under award ECCS-2026822. We thank Noah Eckman for his contributions to GPC data processing.

## TABLE OF CONTENTS

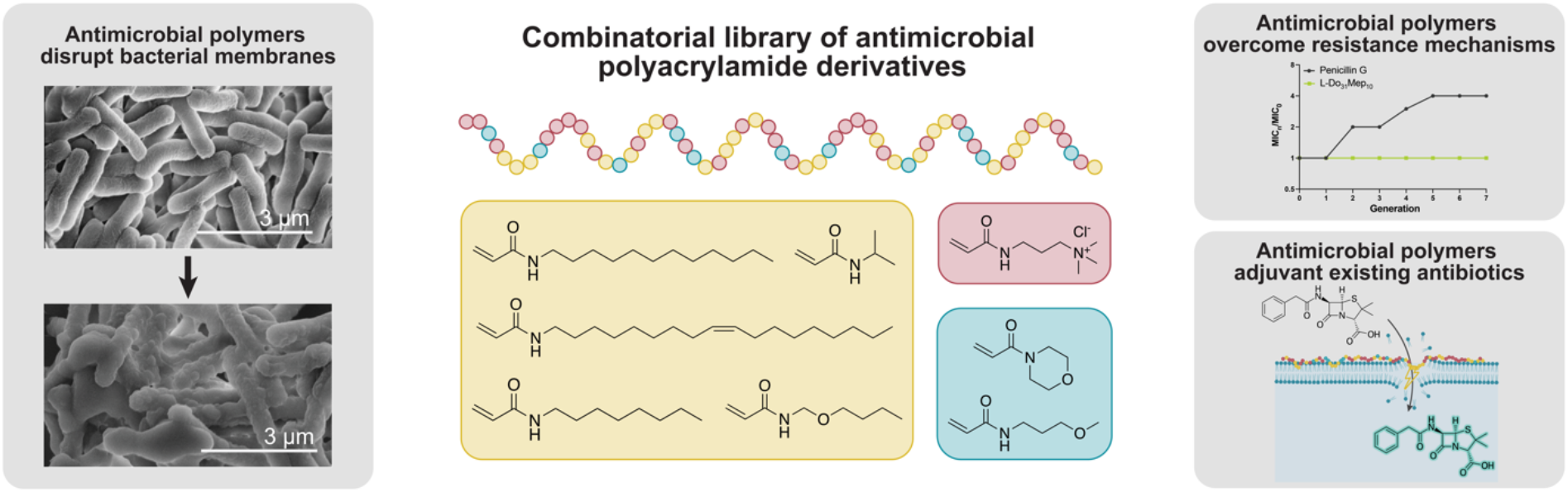

### SYNOPSIS

To combat growing crisis of antibiotic-resistant “superbugs,” we must develop new antibiotics that are effective, safe, and highly stable to facilitate global access. These antibiotics should also overcome typical resistance mechanisms. In this paper, we describe the use of polyacrylamide-based copolymer drugs that function as broad-spectrum antibiotics to combat both Gram-positive and Gram-negative bacteria through a membrane disruption mechanism. These copolymers can delay or prevent the development of resistance and rehabilitate existing antibiotics by improving their potency and limiting the ability of bacteria to develop resistance. To our knowledge, this is the first report of an antimicrobial copolymer that protects a small molecule from the development of resistance. This work represents a significant step towards combatting the antibiotic resistance crisis.

