## Supporting Information for "Polyacrylamide-based Antimicrobial Copolymers to Replace or Rescue Antibiotics"

**Supplemental methods:**

No unexpected safety hazards were encountered.

*Materials*

Solvents *N,N*-dimethylformamide (DMF; >99.7%; Sigma-Aldrich), hexane (>95%, Sigma-Aldrich), acetone (Aldon Corporation), diethyl ether (Fisher Scientific), ethyl acetate (EtOAc; >99.5%, Sigma-Aldrich), methanol (Sigma-Aldrich), deuterium oxide (Sigma-Aldrich), and dimethyl sulfoxide-d6 (Sigma-Aldrich) were used as received. Dichloromethane (DCM; Sigma-Aldrich) was dried over activated molecular sieves before use.

Acryloyl chloride (97%, Sigma-Aldrich), triethylamine (TEA; 99%, Sigma-Aldrich), *N*-octylamine (TCI Chemicals), and oleylamine (Sigma-Aldrich) were used as received.

Monomers *N*-isopropylacrylamide (Ni), *N*-phenylacrylamide (Phe; 99% Sigma-Aldrich), *N*-dodecylacrylamide (Do; TCI Chemicals), *N*-butylacrylamide (Bam; TCI Chemicals), *N*-(Butoxymethyl)acrylamide (Bmam; TCI Chemicals), *N*-(1,1,3,3-Tetramethylbutyl)acrylamide (Tmb; TCI Chemicals) were used as received. 4-acryloylmorpholine (Mo; 97%, Sigma-Aldrich) and *N*-(3-methoxypropyl)acrylamide (Mep; 95%, Sigma-Aldrich) were filtered through basic alumina before use. (3-Acrylamidopropyl)trimethylammonium chloride (Tma; 75 wt. % in $H_2O$, Sigma-Aldrich) was washed thrice with an equal volume of EtOAc and placed under vacuum for two minutes to remove residual EtOAc.

Azobisisobutyronitrile (AIBN; 98%, Sigma-Aldrich) was recrystallized from MeOH and dried under vacuum before use. 4-((((2-Carboxyethyl)thio)carbonothioyl)thio)-4-cyanopentanoic acid (CTA; 95%, Sigma-Aldrich) was used as received.

Bacterial strains were purchased from Microbiologics. Mueller-Hinton media (MHB; BD Difco), Brain Heart Infusion media (BHI; BD BBL), tryptic soy media (Sigma-Aldrich), and agar (Sigma-Aldrich) were autoclaved before use. Cell counting kit-8 (CCK-8) containing the tetrazolium salt WST-8 was purchased from APExBIO. Penicillin G sodium salt was purchased from Research Products International and used as received.

*Synthesis of N-octylacrylamide (Oct)*

*N*-octylacrylamide was prepared by an adaptation of previously described methods.[37] Briefly, *N*-octylamine (992 uL, 6 mmol) and TEA (920 uL, 6.6 mmol) were dissolved in DCM (60 mL). The mixture was cooled in an ice bath. Under a nitrogen atmosphere, acryloyl chloride was added dropwise (536 uL, 6.6 mmol) over a period of 1 h with stirring. The reaction was allowed to proceed for an additional 40 minutes. It was quenched by washing with saturated ammonium chloride (100 mL) followed by saturated sodium bicarbonate (100 mL) and brine (100 mL). The product was dried over anhydrous magnesium sulfate, and the solvent was dried by rotary evaporator. The product (Oct) was isolated as a waxy white solid with 97% yield. NMR spectra were recorded on a Bruker 500 MHz spectrometer.

*Synthesis of oleylacrylamide (Olam)*

Oleylamine (1.974 mL, 6 mmol) and TEA (920 µL, 6.6 mmol) were dissolved in DCM (60 mL). The mixture was cooled in an ice bath. Under a nitrogen atmosphere, acryloyl chloride was added dropwise (536 µL, 6.6 mmol) over a period of 1 h with stirring. The reaction was allowed to proceed for an additional 40 minutes. It was quenched by washing with saturated ammonium chloride (100 mL) followed by saturated sodium bicarbonate (100 mL) and brine (100 mL). The product was dried over anhydrous magnesium sulfate, and the solvent was dried by rotary evaporator. The product (Olam) was isolated as a pale yellow oil with 87% yield. NMR spectra were recorded on a Bruker 500 MHz spectrometer.

*Polymerization conditions*

Statistical copolymerizations were carried out by RAFT polymerization. For low molecular weight polymers, [CTA]/[monomers] = 70. For high molecular weight polymers, [CTA]/[monomers] = 115. [AIBN]/[CTA] = 0.5 for all entries containing the monomer Mep, and [AIBN]/[CTA] = 0.2 for all other entries. The monomers (total mass of 2 g), CTA, and initiator were dissolved in a mixture of DMF and water to a final volume of 8 mL in a 20 mL scintillation vial equipped with a PTFE/silicone septum. The mixture was sparged with $N_2$ for 10 minutes and heated to 65°C overnight. The mixture was cooled to room temperature, exposed to air, and precipitated three times in acetone or a combination of ether and hexane. NMR spectra were recorded on a Bruker 500 MHz spectrometer. $M_n$, $M_w$, and dispersity values were determined via GPC implementing PEG or PMMA standards after passing through an SEC column (Resolve Mixed Bed Low divinylbenzene(DVB) (Jordi Labs)) in a mobile phase of DMF with 1 wt% $LiBF_4$ at 50 °C and a flow rate of 1.0 mL/min (Dionex UltiMate 3000 pump, degasser, and auto-sampler (Thermo Fisher Scientific)).

*Minimum inhibitory concentration (MIC) assay*

The MIC of each polymer was determined by broth microdilution method, adapted from the Clinical and Laboratory Standards Institute (CLSI) guidelines.[38] We used the bacterial strains *E. coli* ATCC 25922, *S. aureus* ATCC 29213, *K. pneumoniae* ATCC 13884, and *E. faecium* ATCC 35667. Briefly, bacteria were cultured in liquid overnight in accordance with ATCC guidelines. *E. coli* ATCC 25922, *S. aureus* ATCC 29213, *K. pneumoniae* ATCC 13884 were grown in MHB, while *E. faecium* ATCC 35667 was grown in BHI. The absorbance of the culture was read at 600 nm, and the culture was diluted to 1 x $10^6$ colony forming units (CFUs)/mL. A two-fold dilution series of 50 µL of polymer solution in media was prepared in a 96-well, U-bottom polypropylene plate. To each well were added 50 µL of the diluted bacterial culture, so each well contained a final concentration of 5 x $10^5$ CFUs/mL. All samples were run in duplicate. Each plate also contained sterile controls (without bacteria) and growth controls (without polymer). The plates were incubated statically at 37 °C for 18-24 h. After incubation, each well was mixed by pipetting to resuspend any bacterial growth, and 75 µL from each well were transferred to a clear, flat-bottom 96-well plate. The absorbance was measured at 600 nm on a

BioTek Synergy H1 microplate reader. The MIC was defined as the lowest concentration before bacterial growth increased by at least 0.1 absorbance units.

*Resistance study*

Development of antimicrobial resistance was monitored by treating bacteria repeatedly with antimicrobial agents. We used *E. coli* ATCC 25922 as a model organism. The same broth microdilution method as used for the MIC assay was employed. After 22-26 h of growth, the well containing ½*MIC was diluted and used to passage the experiment for the next day's growth, so the bacteria were grown continuously in the presence of the antimicrobial agent. For each passage, the MIC was determined by measuring the absorbance at 600 nm, just as for the MIC assay. Resistance was observed by recording changes in the MIC across each passage. All samples were measured in duplicate.

*Membrane integrity study*

Propidium iodide was used to evaluate the integrity of the inner and outer membranes of *E. coli* in the presence of antimicrobial agents, based on previously described approaches.[39,40] Briefly, a stationary phase culture of *E. coli* ATCC 25922 was diluted 1:25 in MHB and grown at 37°C with shaking at 200 rpm to until the absorbance at 600 nm ≈ 0.6. The bacteria were subsequently pelleted by centrifugation at 10,000 rpm for 5 min at room temperature using a Thermo Scientific Sorvall Legend Micro 21R centrifuge. They were washed once with HEPES/glucose buffer (5 mM HEPES and 5 mM glucose). They were pelleted again and resuspended to a final optical density of 0.1 in HEPES/glucose buffer. The cells were aliquoted into a black-walled 96-well plate, and propidium iodide was added to a final concentration of 5 μM. The fluorescence was measured in a microplate for 5 min (excitation 535 nm, emission 617 nm). The plate was removed, and the antimicrobial agents were added at the desired concentrations (from 40x stock solutions prepared in HEPES/glucose buffer). The fluorescence (excitation 535 nm, emission 617 nm) was measured for 1 h at room temperature.

*SEM analysis*

Bacteria were prepared for SEM based on protocols described elsewhere.[32] Briefly, a stationary phase culture of *E. coli* ATCC 25922 was diluted to a concentration of 1 x $10^4$ CFUs/mL in MHB with the desired concentration of antimicrobial agent in a total volume of 1 mL. The mixture was incubated statically at 37°C for 2 h. Bacteria were collected by centrifugation at 4,000 xg for 5 min at room temperature and washed twice with phosphate-buffered saline (PBS). The samples were fixed in 800 μL of a 5% formaldehyde solution at room temperature for 1.5 h. The samples were pelleted by centrifugation at 4,000 xg for 5 min and washed thrice with water. The samples were dehydrated by graded ethanol solutions (35% for 5 min, 50% for 5 min, 75% for 5 min, 90% for 10 min, 100% for 10 min). The samples were resuspended in a minimal volume of ethanol (~15 μL) and drop-cast onto a copper 300 mesh grid, secured with silver paste to a flat aluminum pin stub. A 3.5 (±0.5) nm thick layer of pure gold was deposited onto the samples using the Leica ACE600 Vacuum System. SEM analysis was conducted using the FEI Magellan 400 XHR Scanning Electron Microscope, operated under high vacuum at 5.00 kV, 25 pA with beam deceleration in immersion mode.

*Hemolysis assay*

Red blood cells (RBCs) were collected by centrifugation of whole blood at 500 xg for 5 min. The plasma was removed, and the RBCs were washed twice with 150 mM sodium chloride and once with PBS. They were pelleted again and resuspended in PBS. The sample was diluted 1:50 in PBS. To each well of a 96-well plate was added 180 µL of diluted RBCs and 20 µL of antimicrobial agent stock solution at 10x the desired concentration. Samples were run in duplicate. Triton X-100 at a final concentration of 1% was used as a positive control, and water was used as a negative control. The plate was incubated statically at 37$^{o}$C for 1 h. Intact RBCs were pelleted by centrifugation at 1,000 xg for 5 min using a Thermo Scientific Sorvall Legend XTR centrifuge. 100 µL of the supernatant were transferred from each well to a clear, flat-bottom 96-well plate, and the absorbance was measured at 450 nm. The percentage of hemolysis was calculated as follows:

Hemolysis (%) = [($Abs_{450\ nm}$ of the treated sample – $Abs_{450\ nm}$ of the negative control)/($Abs_{450\ nm}$ of positive control – $Abs_{450\ nm}$ of negative control)] × 100%.

*Cytotoxicity assay*

Cellular cytotoxicity was evaluated *in vitro* using a CCK-8 colorimetric kit to quantify the viability of 3T3 cells and A459 cells. Briefly, cells were plated at a concentration of 10,000 cells per well in a clear, flat-bottom 96-well plate. The cells were allowed to adhere overnight. The antimicrobial agents were dissolved in media and added to each well at the desired concentrations to reach a final volume of 100 µL. Each sample was run in triplicate. A negative control (no cells) and an untreated sample (no polymer) was included on each plate. After 5 h of incubation with polymer, 8 µL of CCK-8 solution was added to each well, and the plate was returned to the incubator. 24 h after the addition of polymer, the absorbance was measured at 450 nm. The percentage of viability was determined as follows:

Viability (%) = [($Abs_{450\ nm}$ of the treated sample – $Abs_{450\ nm}$ of the negative control)/($Abs_{450\ nm}$ of untreated sample – $Abs_{450\ nm}$ of negative control)] × 100%.

The LC50 was determined through interpolation after fitting the data to a five-parameter asymmetric sigmoidal function using GraphPad Prism software.

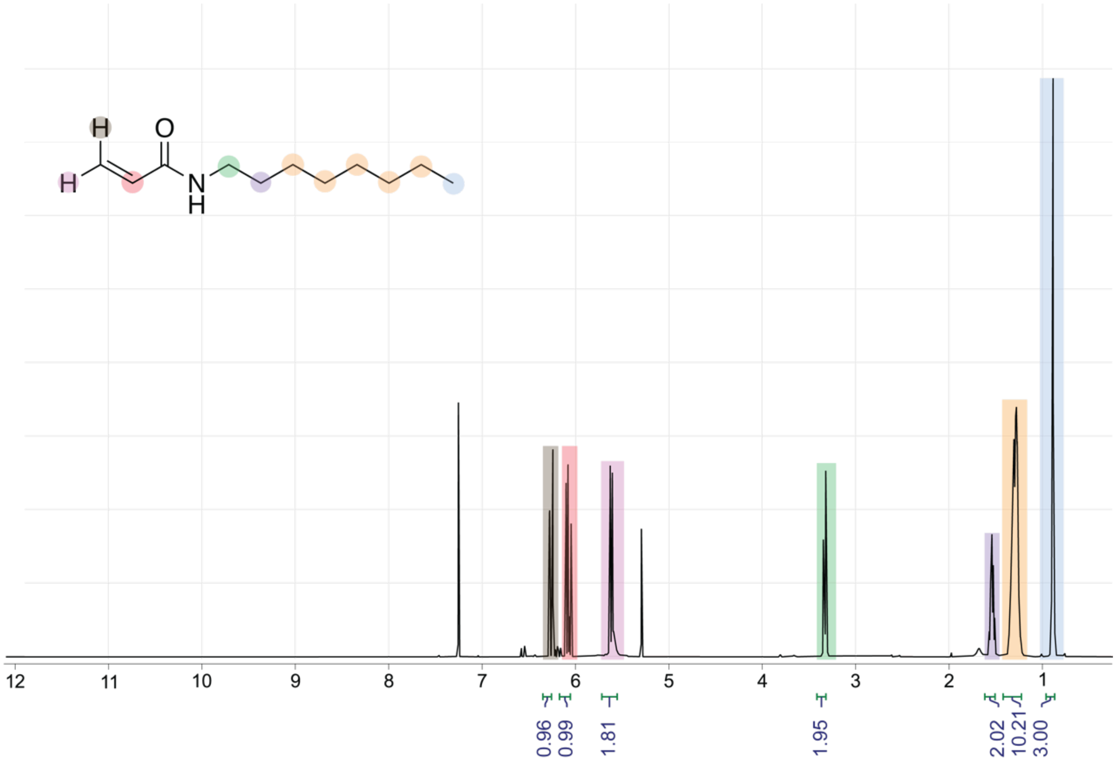

**Figure S1. NMR Spectrum of Octylacrylamide.** Measured in $CDCl_3$ using a 500 MHz NMR magnet.

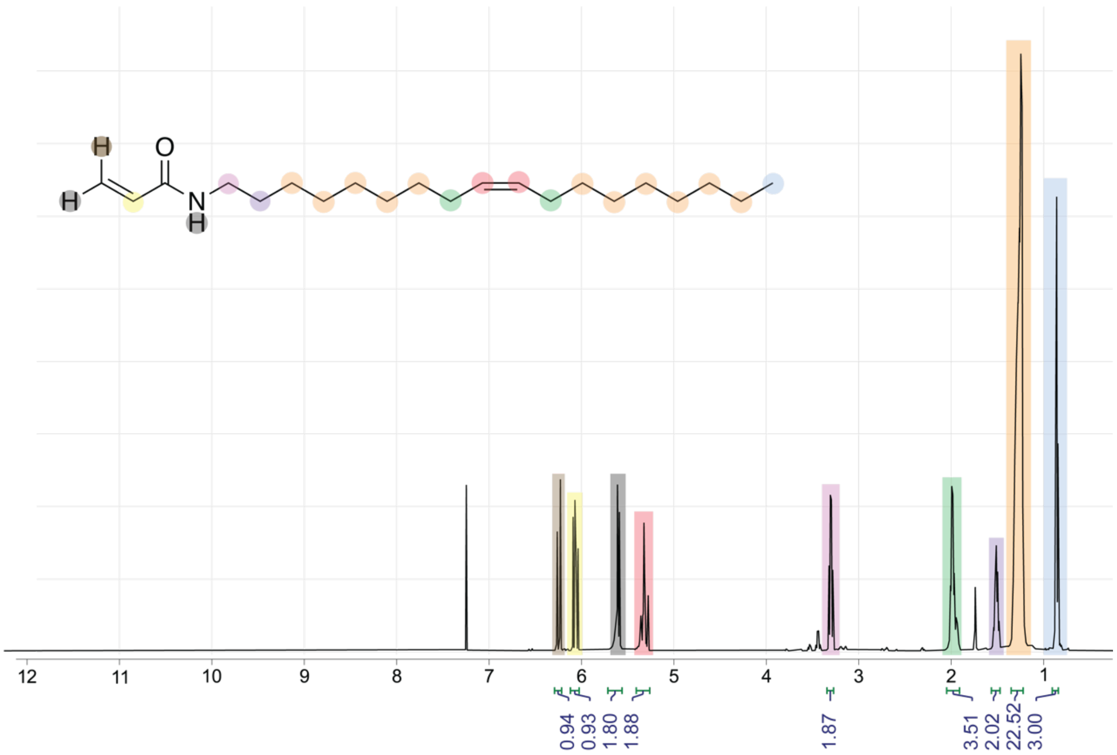

**Figure S2. NMR Spectrum of Oleylacrylamide.** Measured in $CDCl_3$ using a 500 MHz NMR magnet.

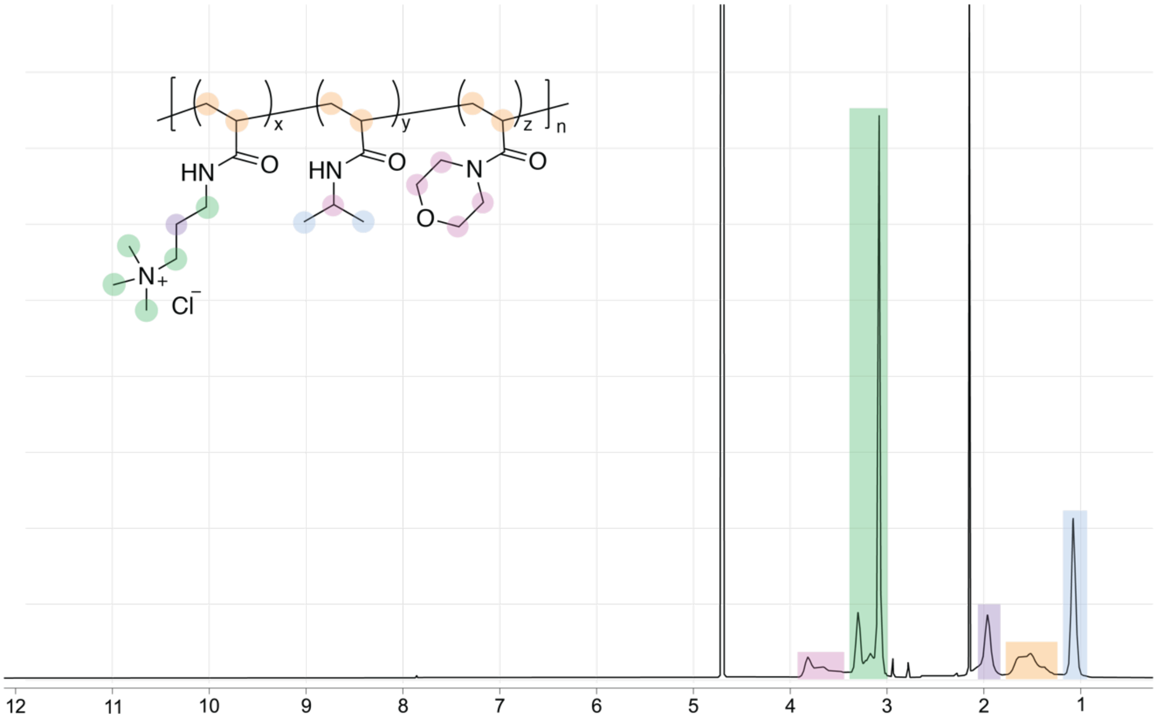

**Figure S3. NMR Spectrum of L-Ni$_{31}$Mep$_{10}$.** Measured in $D_2O$ using a 600 MHz NMR magnet.

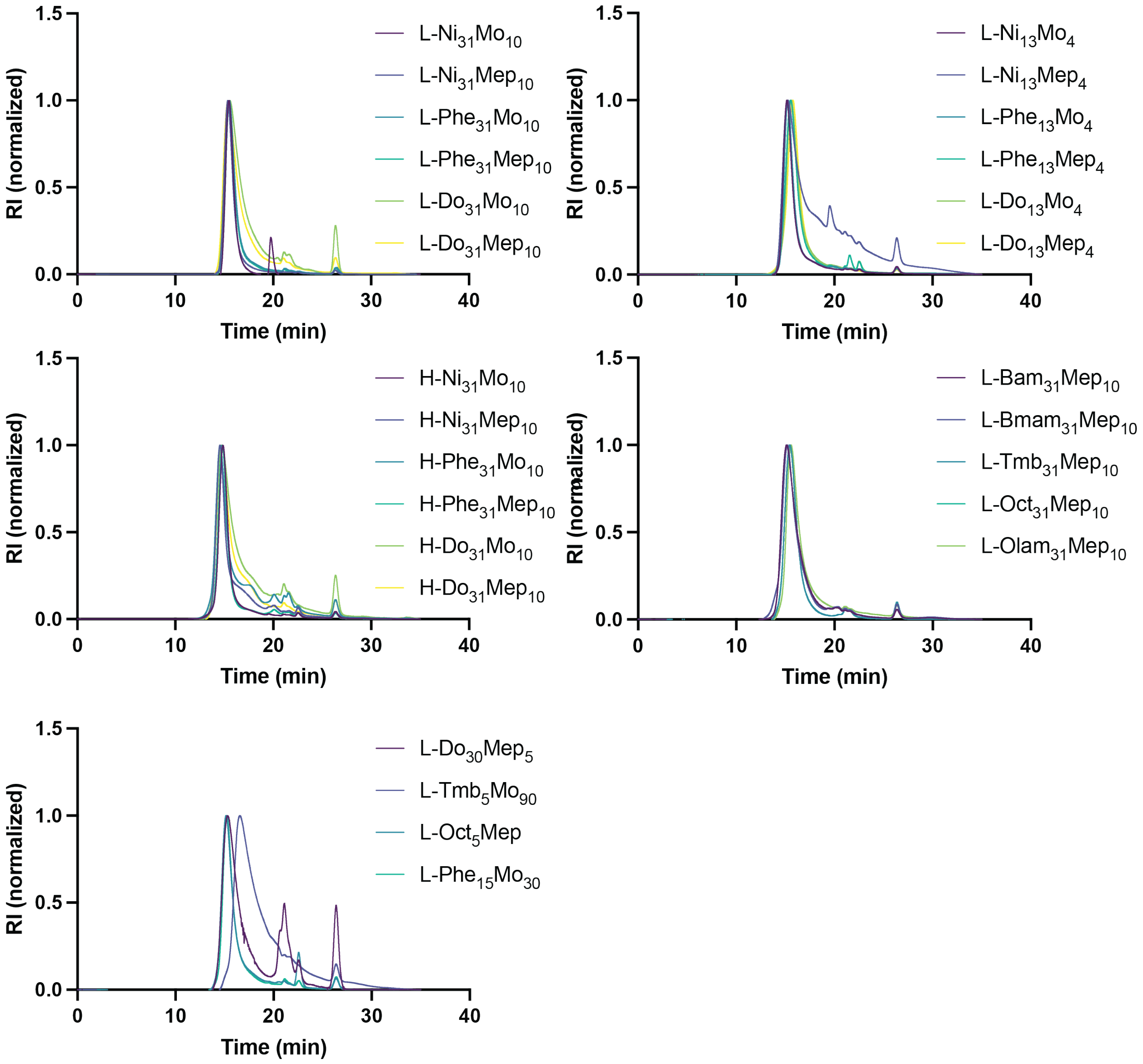

**Figure S4. GPC traces.** Normalized and baseline-corrected.

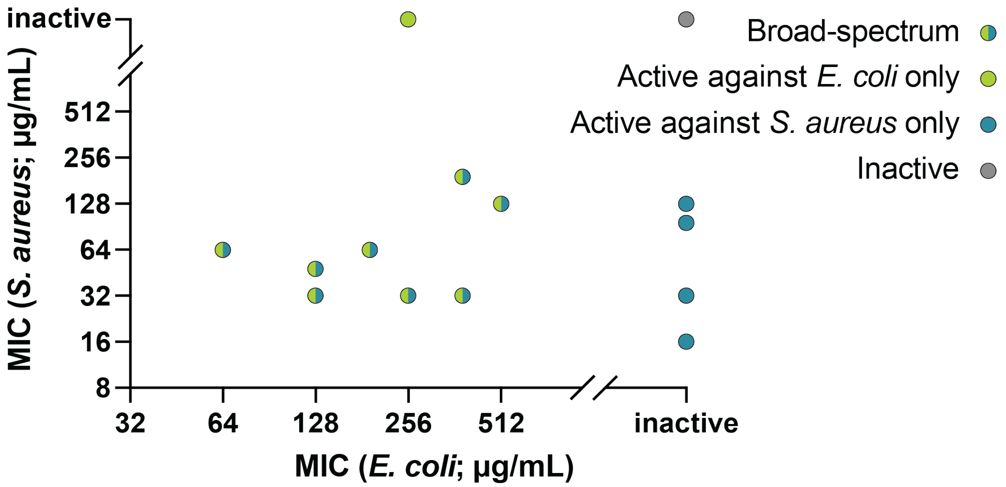

**Figure S5. Broad antimicrobial activity of polymer library.** Each dot represents a novel copolymer, where its location is defined by its efficacy against *S. aureus* and *E. coli*.

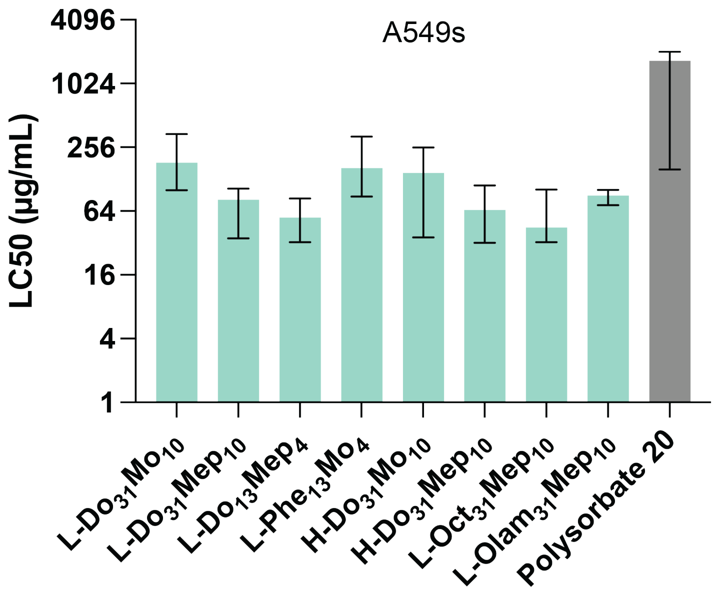

**Figure S6. Cytotoxicity of top-performing copolymers in A549 cells.**

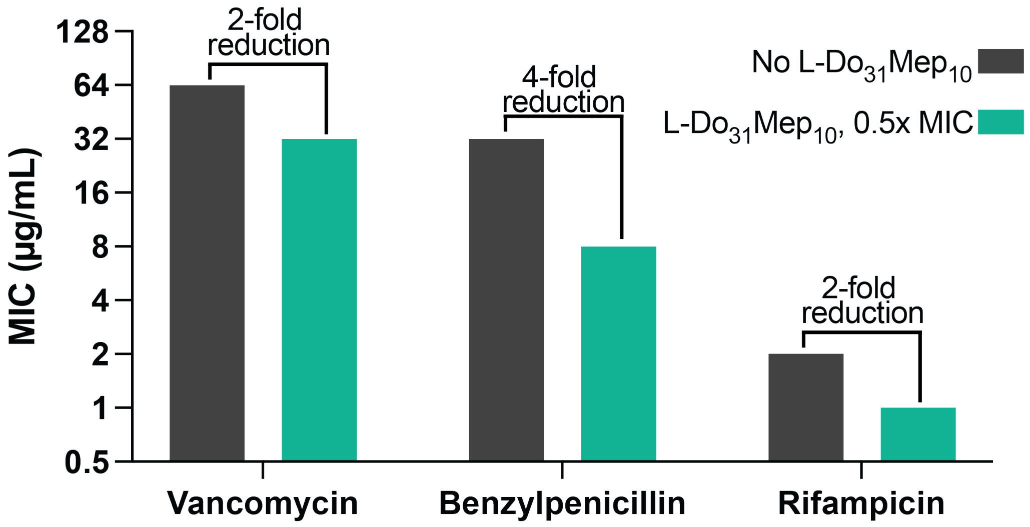

**Figure S7. Adjuvanting efficacy of L-$Do_{31}Mep_{10}$.** The presence of L-$Do_{31}Mep_{10}$ reduces the MIC of vancomycin, penicillin, and rifampicin, suggesting that their effects are additive.
